# Bumblebee visual learning: simple solutions for complex stimuli

**DOI:** 10.1101/2024.03.16.585132

**Authors:** Marie-Geneviève Guiraud, Vince Gallo, Emily Quinsal-Keel, HaDi MaBouDi

## Abstract

Natural visual stimuli are typically complex. This presents animals with the challenge of learning the most informative aspects of these stimuli while not being confused by variable elements. How animals might do this remains unclear. Here, we tested bumblebees’ ability to learn multicomponent visual stimuli composed of a simple constant bar element and a grating element that was consistent in orientation but varied in width and number of gratings. Bees rapidly and successfully learned these compound stimuli. Tests revealed learning of the single bar element was more robust than learning of the grating element. Our study highlights how even small-brained invertebrates can rapidly learn multicomponent stimuli and prioritise the most consistent elements within them. We discuss how the learning phenomena of generalisation and overshadowing may be sufficient to explain these findings, and caution that complex cognitive concepts are not necessary to explain the learning of complex stimuli.

**Highlights:**

- Bumblebees are highly efficient in prioritising the most consistent elements in multicomponent visual stimuli.
- Bees trained on horizontal and vertical cues exhibit differences in how they memorise visual cues.
- Two phenomena can explain how bees preferentially select, memorise and use visual cues in this experiment: generalisation and overshadowing.
- Bumblebees as generalist foragers are well-suited to study visual cognition.

## Introduction

Natural stimuli are rarely simple. Flowers, for instance, are multimodal stimuli, and even within just the visual domain flowers vary in colour, size, structure, and luminance. In this study, we challenged bumblebees in a learning assay with multicomponent visual stimuli to explore how bees learn complex stimuli.

For this work, we adopted a definition of visual complexity from computer vision where complexity encompasses order (repetition and redundancy), variety (Tatarkiewicz 1972, Tsotsos 1990, Simoncelli et al. 2001), compactness, as well as the numbers of lines and edges of varied orientations, and open and closed figures in an image (Biederman 1987, García, Badre et al. 1994, Mirmehdi, Palmer et al. 1999). Complex images typically contain a greater number of edges and less predictable distribution of edges across the images, whereas simple images contain redundant and predictable data and are, therefore more compressible (Tatarkiewicz 1972, Tsotsos 1990). Both humans and computer learning algorithms recognize simple elements present within complex images to facilitate visual learning (Rahardja 1996, Biederman 198, Szeliski 2022), but it is less clear how animals might learn complex visual stimuli.

For this question, the bee is an excellent model as a very good visual learner, they can rapidly learn associations between visual stimuli and punishment or reward (Avargues-Weber et al. 2011, Guiraud et al. 2022). They excel in object recognition and have the capacity to generalize what has been learnt to similar stimuli thereby creating categories of objects (Gould 1985, Hateren, Srinivasan et al. 1990, Horridge 2000, Srinivasan 2010, Avargues-Weber, Deisig et al. 2011). In the visual domain, bees can learn to discriminate items very rapidly, even recognise human faces (Dyer, Neumeyer et al. 2005) and prefer global visual cues over local visual cues (Avargues-Weber et al. 2015). Moreover, bees have been shown to recognise classes of objects by shared “abstract” properties like relative position (above / below for instance: Avargues-Weber et al. 2011, Guiraud et al. 2018) or relative size (Avargues-Weber et al. 2014).

Prior studies have suggested that when learning complex visual stimuli bees selectively attend to discrete aspects of visual information and ignore irrelevant perceivable surrounding information (Spaethe, Tautz et al. 2006). For example, bees can select a rewarding configuration of oriented bars over a variable, distractor pattern with the same orientation (Stach and Giurfa 2005). This suggests that bees have an ability to focus on the most salient visual cue present during training. Moreover, the length of training appears to modulate this attention (Stach and Giurfa 2005). Presently, we do not know what cognitive abilities might allow bees to selectively attend to the most salient part of a complex stimulus. To explore this issue, we tested bumblebees’ learning of visual stimuli that contained two elements. One element was simple and constant (either a vertical or horizontal bar). One was more complex and variable (gratings of constant orientation but variable sizes, number of bars and widths). During tests, we examined what elements of these complex stimuli had been learned by the bees, and how well they might generalise the learned stimuli.

## Material and methods

Forty-one bumblebees (*Bombus terrestris audax)* from nine commercially available colonies (Agralan Ltd., Swindon, UK) were trained and tested. Each colony was maintained in a wooden nest box (28 cm L× 16 cm W × 11 cm H). This was connected to a Perspex tunnel (1.5cm^2^ and 20cm long) leading to a flight arena (60cm l × 60cm L × 40 cm H), in which workers could freely forage for 30% sucrose solution (w/w) during non-training periods. Pollen was provided to the colony every two days. The arena was covered by ultraviolet-transparent Plexiglas. The walls of the flight arena were covered with a laminated pink and white Gaussian random dot pattern to provide optic flow for the bees and contrast for video recording. High-frequency fluorescent lighting mimicking natural light (containing both UV and the full spectrum of visible light: TMS 24F lamps with HF-B 236 TLD ballasts, Phillips, Netherland; fitted with Activa daylight fluorescent tubes, Osram, Germany) illuminated the arena (Fig. 1).

**Figure 1.**
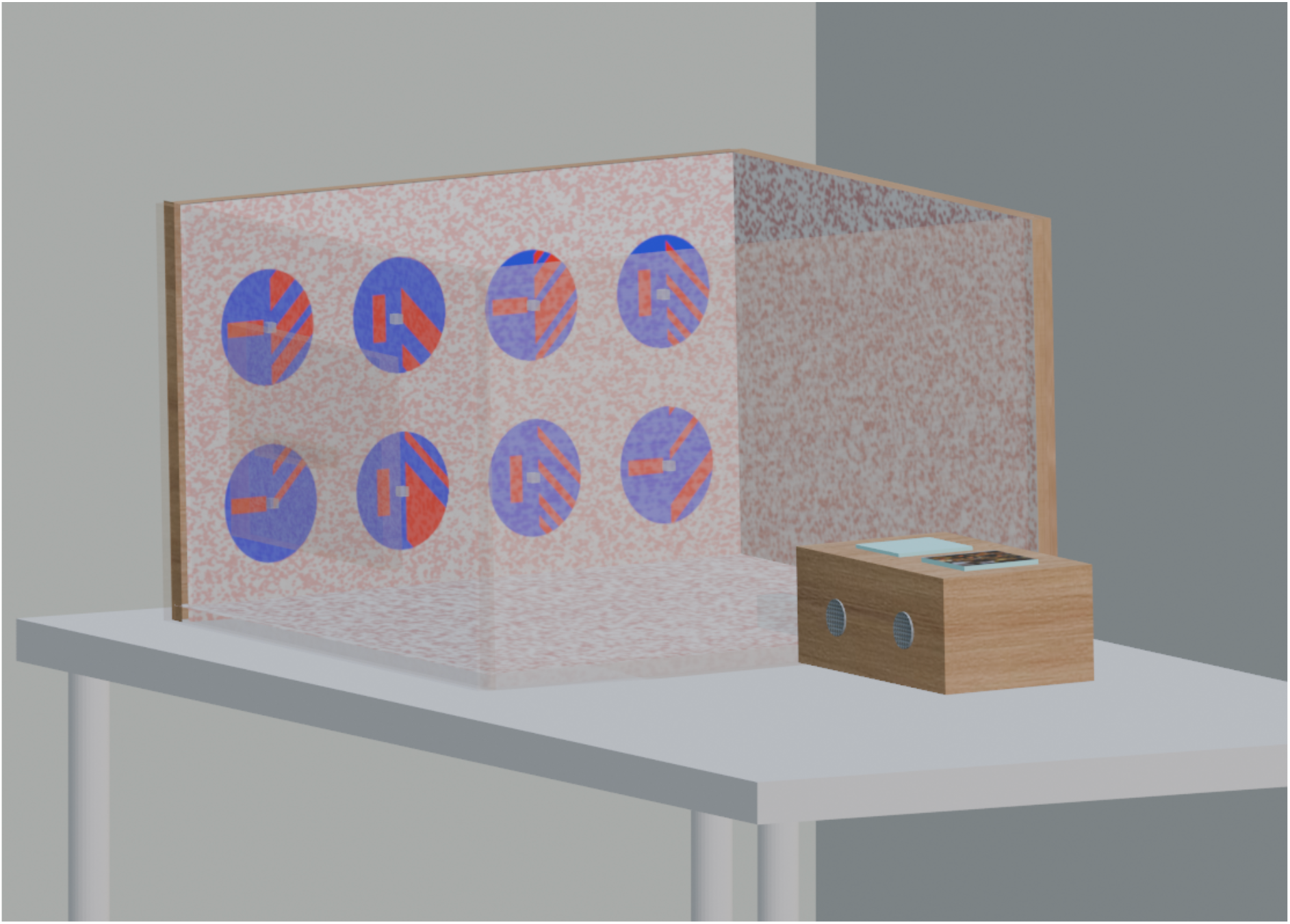
Flight arena. The back if the arena displays eight stimuli with a feeding station. Half of the stimuli (CS+) provided sucrose solution and the other half (CS-) provided quinine solution (aversive). Positions and stimuli varied between trials.

During training, one individual worker could forage from eight feeding stations. The feedings stations were transparent concave Perspex cubes (1,5 cm2 and 0.8 cm high with a hole ⌀=0.6cm, depth 0.3cm; see Fig. 1). They were positioned vertically (using blue tack) onto the experimental wall presenting the visual stimuli. Each feeding station delivered 15 μl 50% sucrose solution (w/w). The small volume of sugar solution (15 μL) delivered by each feeding station was well under the crop capacity of bumblebee, which encouraged the bee to visit multiple feeders in a foraging trip.

During pre-training, the selected worker would be familiarised with drinking from all the feeding stations. Workers successfully using the feeders were marked with coloured number tags (Opalithplättchen, Warnholz & Bienenvoigt, Ellerau, Germany).

### Visual stimuli

Visual stimuli (Fig. 1) were printed using a high-resolution colour laser printer. Stimuli were covered with transparent film, allowing them to be cleaned with 70% ethanol after every trial to remove odours and pheromones potentially left by bees. The background of all stimuli was always a blue (RGB colour 0,0,255) 8.5 cm diameter circle on which was printed a red (RGB colour 255,0,0) design. Each stimulus contained multiple elements; therefore, more than one element was associated with the CS+ during training. One half of the stimulus was either a vertical or horizonal bar (width: 12.0 mm, length 35.0 mm, see Fig. 1). The other half of the stimulus was a grating oriented at either +45 ° or −45 °. The grating widths varied from 7.0 mm to 57.0 mm and extended from the vertical centre of the circle to its outer perimeter. These visual cues were randomly selected patterns generated by MATLAB. (version 2015b; The MathWorks, Inc., Natick, MA, USA), from a set where the number, size and spacing of the bars varied (Supplementary Fig. S1). In total, 15 versions of each of the ‘vertical’ and ‘horizontal’ stimuli were created (Fig. S1). During training, both the orientation of the bar (vertical or horizontal) and the orientation of the gratings (+ or – 45 °) were reliably associated with the CS+.

Four stimulus groups were defined for training based on the rewarding stimuli used (Fig. 2). The H1 group (10 bees) had for CS+ stimulus a horizontal bar on the left and −45 ° gratings on the right. The H2 group (10 bees) was the mirror image of this (a horizontal bar on the right and 45 ° gratings on the left). The V1 group consisted of 11 bees with the CS+ stimulus a vertical bar on the left and −45 ° gratings on the right. For the V2 group (10 bees), the CS+ was the mirror image of V1. In each group, the CS-was the opposite of the CS+ (Fig. 2). Comparing the four groups allowed us to test if placement of elements within the stimuli influenced the results. CS+ was associated with 15 μl 50% sucrose solution (w/w), while CS-was presented with saturated quinine hemisulfate solution (15 μl).

**Figure 2.**
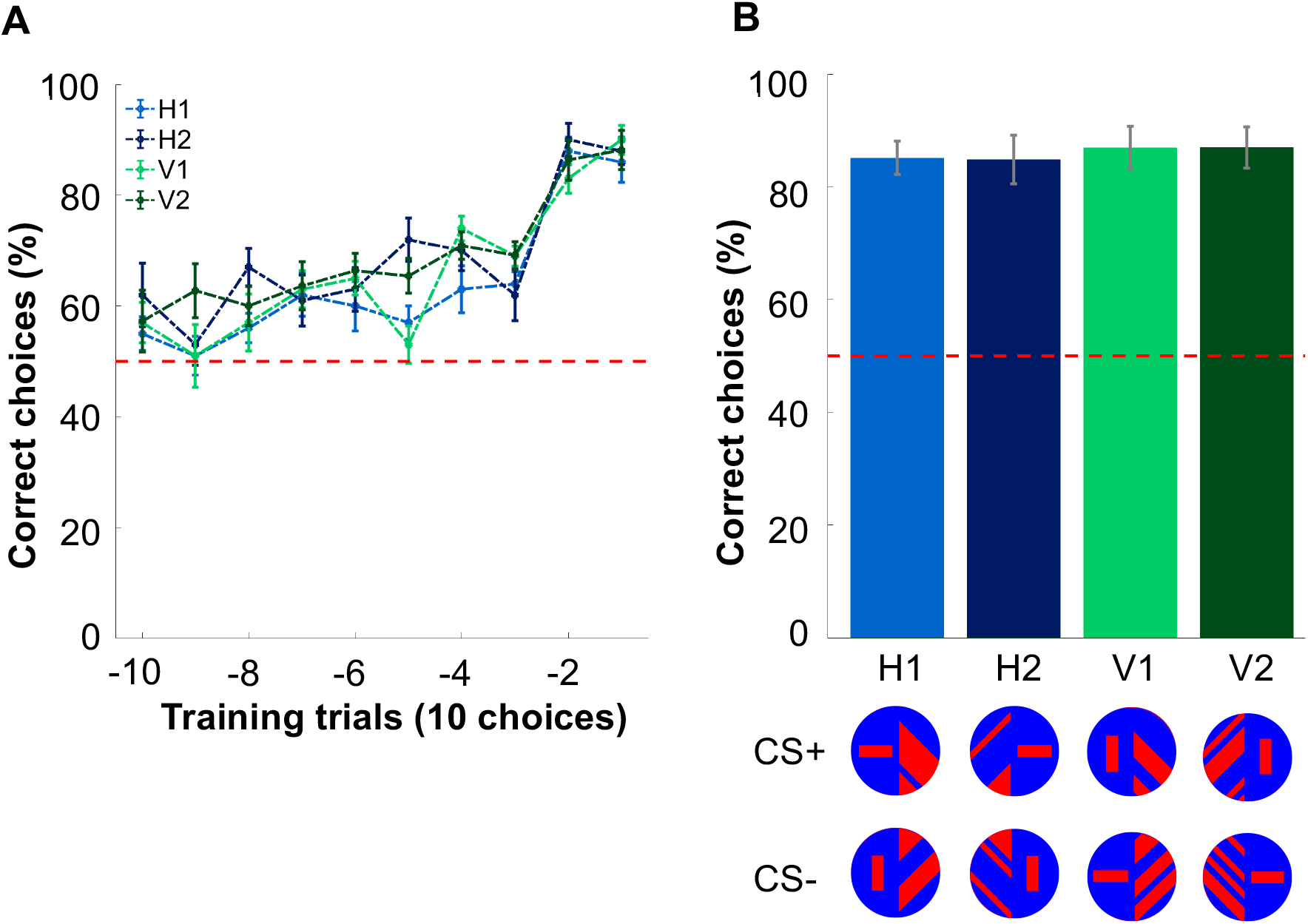
Performances of bees during the training and the learning test. A. The mean percentage and standard error of the last 100 choices by the bees are plotted as a function of blocks of 10 choices for the four training groups (mean ± SEM) Training concluded when a bee reached the performance threshold (>80% correct choices over 20 trials) hence all bees made a different number of choices during training. Here we plotted the last 100 choices (by blocks of 10 choices) and the x axis counts these blocks down to threshold (block −10 to 0). **B. Percentage of correct choices of each bee in the unrewarded learning test for the four training groups (mean ± SEM).**

For the tests, three other types of stimuli were produced (Supplementary Fig. S1). In the Conflict test, for each group the test stimuli swapped the orientation of gratings between the CS+ and CS-so that the presented stimuli now contained elements of both CS+ and CS-(Fig. 3). Half pattern tests presented stimuli that only contained some of the elements or the complex stimuli: only bars or only gratings appearing on the blue background (Fig. 3). Of the eight stimuli presented to bees in the test, bees were presented with two stimuli with only a horizontal bar, two stimuli with only the vertical bar, two stimuli with +45° gratings and two stimuli with −45° gratings. This tested which elements of the compound stimuli had been learned and were most preferred by trained bees. For the generalisation test, the same pattern configurations as in the training stimuli groups was used, except the horizontal (‘H1’, ‘H2’) or vertical (‘V1’, ‘V2’) bars were replaced with two parallel bars (the original one: width: 12 mm, length 35 mm; and the second with width 5 mm to 8 mm, length 35 mm, with a 11 mm separation) centred within the respective pattern halves (Fig. 4). This tested whether bees could generalise learning a complex stimulus to a similar stimulus. There were eight stimuli shown in each test.

**Figure 3.**
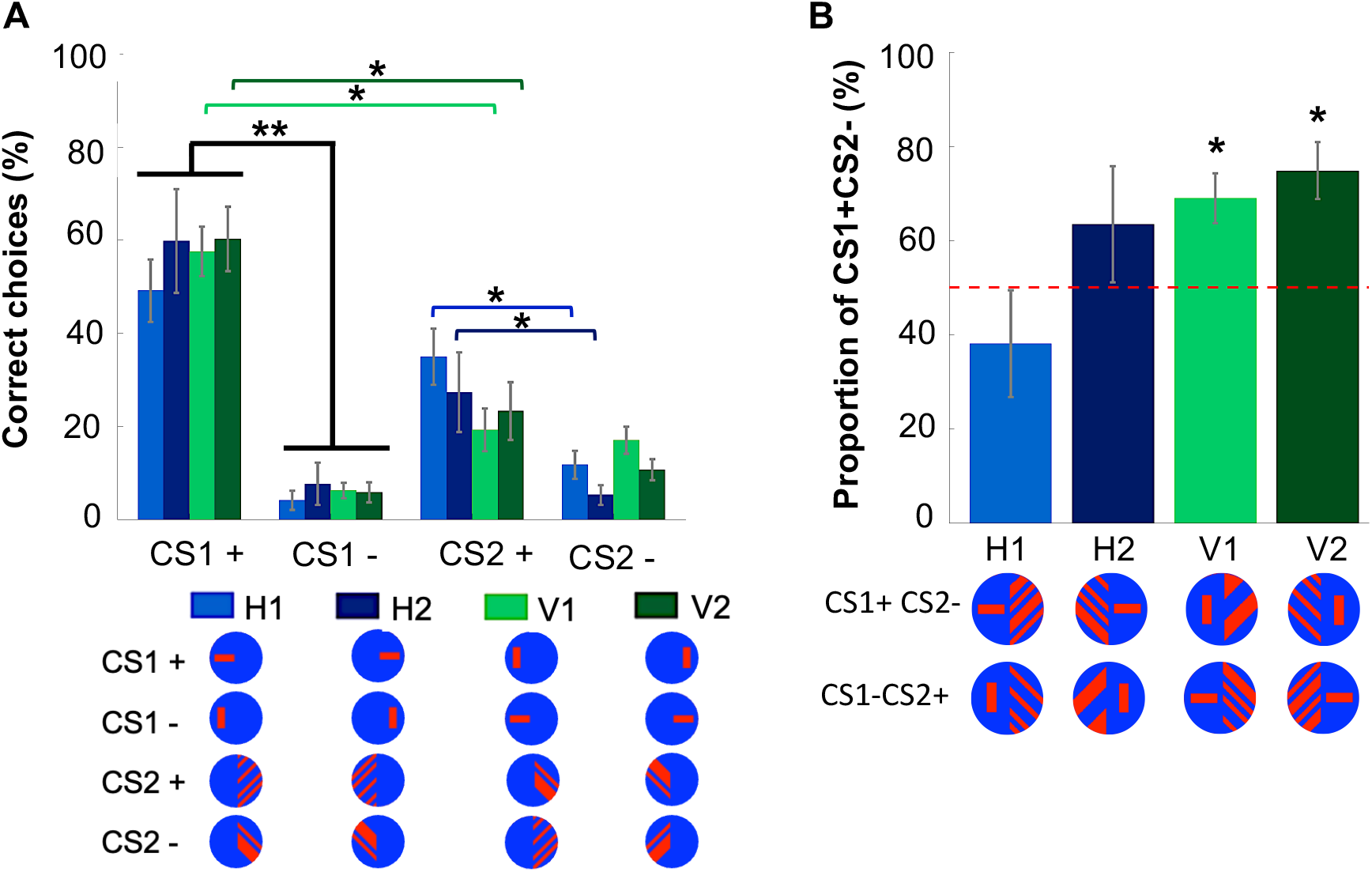
A. Percentage of choices by each bee for each stimulus element in the half pattern test. Bars indicate training groups. For each group (H1, H2, V1, V2) the correct CS1+ (part 1 of the conditioned stimulus) and the correct CS2+ (part 2 of the conditioned stimulus) are to be found in Supplementary Table 1 (mean ± SEM). **B. Percentage of CS1+CS2-choices by each bee in the four groups during the conflict test.** Only bees trained on the vertical stimuli choose the CS1+CS2-above chance level (red dotted line) (mean ± SEM). **p<0.05, **p<0.01 and ***p<0.001 vs. random choice.

**Figure 4.**
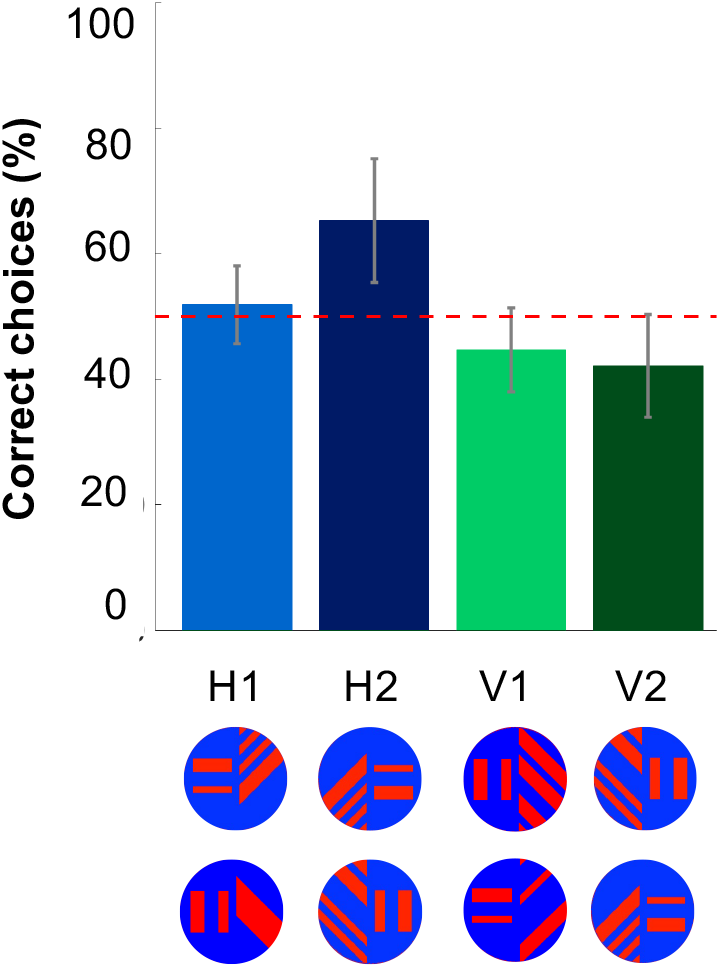
Percentages of correct choices by each bee from the four groups during the generalisation test. Data are presented as the mean ± S.E.M.

### Training and tests

Pre-training encouraged the bee to visit each of the feeder locations. For this, all the stimuli were plain blue disks and all of them provided 15μl of 50% sucrose solution (w/w).

After pre-training, the flight arena was emptied of the bees and thoroughly cleaned with 70% ethanol to remove potential olfactory cues. A selected bee was assigned to one of the four stimulus groups (H1, H2, V1, V2). In each trial, four of the fifteen available pattern variations, for each stimulus group, were pseudo-randomly selected. Each pattern did not appear more than once in consecutive bouts. Eight stimuli were shown in each trail: four CS+ and four CS-. These were pseudo-randomly placed on the presentation wall to prevent the bees establishing location biases (Fig. 1). CS+ stimuli were replenished with sucrose solution once the bee had landed on all rewarding feeders.

In each trial, a choice was considered as each landing a bee made on the feeding stations in the trial. The number of choices usually varied between 8 and 12 before the bee went back to the nest. For consistency, and only for the training trials (Fig. 2A), we plotted the bee choices by blocks of 10 choices. The training phase ended when a bee exhibited > 80% CS+ choices in the last twenty choices. Due to the nature of this threshold the number of training trials and choices varied between 90 and 180 choices made, with on average 140 choices made before a bee reached the threshold. Individual bee training took between 4 and 8 hours to be completed. Three bumblebees failed to complete the training phase (they did not return to the flight arena during the training period) and are not included in these data. For all tests, all choices are accounted for during a period of 2 minutes.

Following the last training trial, the non-rewarded tests began. During tests, all stimuli offered only 15 μl distilled water. Both the place of stimuli and the sequence of tests were randomized between bees. Bees were exposed to refreshment trials in between tests using the reinforced training stimuli and their performance was assessed. Bees progressed to the next test once they achieved > 80% correct choices over 20 consecutive choices. We gave each bee four tests. (1) a learning test with novel stimuli (with similar configurations as the training patterns); (2) a conflict test with conflicting stimuli; (3) half-pattern test presented only one side of the patterns, single bar or gratings alone, was presented to bees, from both the rewarding (CS+ 1 and CS+ 2) and aversive pattern stimuli (CS-1 and CS-2). Finally, (4) a generalisation test consisted of similar pattern configurations of that used during training.

### Data analysis

Data from the training trials were analysed using a logistic regression *via* a Generalised Linear Mixed Model (GLMM), which evaluated the performance of the four groups of bees. The performance of a bee throughout the training procedure was calculated as the percentage of correct choices for every consecutive block of 10 choices (Fig. 2 for the last 10 trials / 100 choices). The blocks of 10 choices, the four training groups of bees and the interaction between the choice block and the training groups were considered as explanatory variables in the model. Finally, the GLMM’s parameters were estimated by Maximum likelihood method in MATLAB (2022b).

To assess whether performance differed between the four groups during the tests, non-parametric statistical tests were used. In the tests, for each bee, we calculated the percentage of correct choices (CS+ stimulus) in a two-minute period. A Kruskal-Wallis test was used to determine whether there was any difference between the training groups of bees when they were confronted with novel stimuli in the learning and generalisation test. A Wilcoxon signed-rank test was applied to all tests to compare the performance of bees against the performance level expected by chance, and to identify visual cue (bars or gratings) preference in individuals (half-pattern test). The same test was used to compare the responses of bees to test stimuli where H1 and H2 were pooled to form the H group, and V1 and V2 were pooled to form the V group. All statistical analyses were conducted with MATLAB (MathWorks, 2022b). Data is presented in figures using SEM.

## Results

Bees from all groups significantly increased their number of correct choices over time (GLMM P=1.407×10^−43^, Table 1, Fig. 2A). No significant difference in the proportion of correct choices between groups of bees was found (GLMM, P=0.105; Table 1).

**Table 1.** Summary of the Generalised Linear Mixed Model (GLMM) examining factors that contribute to variation in performance during training. The dependent variable was the number of correct choices from the block of 10 choices. Fixed factors such as group, beegroupHV, and trial were examined in the model. Bee index was considered in the model as a random factor. Formula: response ∼ 1 + trials + beegroupHV + beegroup + (1 | bee index). Model fit statistics: AIC= 1077.5, BIC= 1099.2, LogLikelihood= −533.76, Deviance=1067.5. This model is the best model with lower BIC value that provides a better trade-off between fit and complexity (number of parameters). The length of the training until bees reached the training criteria was not statistically different between groups (Supplementary table 1A. Kruskal-Willis test df=39, Chi-sq=2.47, p=0.48).

|  | Estimate | SE | tStat | pValue |
| --- | --- | --- | --- | --- |
| **Intercept** | -0.150 | 0.125 | -1.197 | 0.231 |
| **Group index2**<br>**("H" or "V")** | -0.235 | 0.158 | -1/486 | 0.137 |
| **Bee group** | 0.113 | 0.069 | 1.622 | 0.105 |
| **Trials** | 0.099 | 0.006 | 15.104 | 1.407 e-43 |

Bees from all groups successfully recalled the association of the visual stimulus with the sucrose reward as they performed above 80% correct choices on average during the learning test (Supplementary Table 1B. difference to chance level: Wilcoxon signed rank test p<0.05 for each group, Fig. 2B). No difference in the performance of bees between groups was observed (Supplementary Table 1B. Kruskal-Wallis test: df=39, Chi-sq=2.47, p=0.48, Fig. 2B).

### Half-pattern test

The half-pattern test examined what elements of the visual stimuli the bees had learned (Fig. 3A). In all four training groups, bees clearly learned the orientation of the simple bar stimulus, showing a strong preference for the rewarded orientation over the punished orientation (Fig. 3A). The gratings element was learned less well. Only bees in the H groups (where CS+ was associated with the horizontal single bar and related gratings) preferred the rewarded grating orientation to the punished grating orientation. Bees in the V groups (where CS+ was associated with the vertical single bar and related gratings) did not differ in their choice of rewarded or punished grating (Supplementary Table 2). Bees in all groups chose the rewarded bar more than the rewarded grating, but this difference was only significant for bees in the V groups (Fig. 3A, Supplementary Table 2).

### Conflict test

In the conflict test, stimuli combined elements of the rewarded and unrewarded stimuli (Fig. 3B). Similarly, to the ‘half-pattern’ test, only bees trained on the pattern containing a single vertical bar and 45° gratings (V1 and V2) choose more often the conflicting pattern containing the single vertical bar over the other one (Supplementary Table 1). Bees trained on the pattern containing the single horizontal bar choose both conflicting patterns equally (Supplementary Table 3). No difference in performance was found depending on each groups’ cues side (H1 and H2, V1 and V2).

### Generalisation test

No group of bees were able to generalise the trained stimuli to a new stimulus with two bars, and no difference was found between groups (Fig. 4, Supplementary Table 4).

## Discussion

In our study, all bees were able to learn the complex visual stimulus (Fig. 2), but bees learned the simple bar element of the complex visual stimulus better than the grating element (Fig. 3). Following training on the compound stimulus, all groups of bees learned to prefer the orientation of the single bar over the oriented gratings (Fig. 3). For the gratings, only bees in the groups where the CS was associated to the horizonal single bar and the 45° gratings (H groups) showed a significant preference for the rewarded orientation over the punished orientation. This can be partly explained by their scanning behaviour when approaching these patterns (MaBouDi et al. 2021), but also due to the nature of their visual system (Guiraud et al. 2023). All four groups chose bar elements more than grating elements, but this difference was only significant for bees from the group that associated the CS+ with the vertical single bar and the 45° gratings (V groups, Fig. 3A).

In our experiment, the bar element was simpler than the grating element in that it had fewer lines and edges (according to computer vision definition, Szeliski et al. 2022). It was also less variable since it did not vary in shape or position during training. By contrast, grating elements had a constant orientation but the number, width and spacing of the gratings varied during training trials. In our training, bees had to simply learn to discriminate the horizontal and vertical bars, whereas for the gratings they had to learn to discriminate categories of +45° and −45° gratings. Learning to discriminate categories of stimuli is slower than learning to discriminate individual stimuli in bees and other animals (Zhang et al. 2004, Wehner 1967, 1971, Bernard et al. 2006 Stach et al. 2004, 2005). This difference likely contributed to the stronger learning of the single bar seen in our data.

If the bar element was learned faster than the grating element in training, then the bar may have overshadowed learning of the grating. Overshadowing is a well-established learning phenomenon in many animals. It describes a phenomenon where if an animal is conditioned with a compound stimulus AB, the subsequent response to B would be less than if it had received a similar amount of training with B alone (Brembs & Heisenberg 2001, Linster & Smith 1997, Pavlov I.P. 1927). Overshadowing can be asymmetric, with the most salient element of the compound stimulus more likely to overshadow the less salient (Smith et al. 1994).

In bees, overshadowing has been demonstrated in olfactory conditioning (Linster & Smith 1997, Pelz et al. 1997, Schubert et al. 2015). Linster & Smith (1997) have argued that it is not necessary to invoke attentional and higher-order cognitive processes to explain overshadowing. They have proposed a model that can explain the overshadowing phenomenon as a result of processing between the olfactory glomeruli and reinforcement neurons within the antennal lobe. Overshadowing is considered a key element of many fundamental learning theories, and it has been demonstrated in visual and olfactory learning domains in many vertebrates (Mackintosh 1971, Tennant et al. 1975, Sherratt et al. 2015). Brembs & Heisenberg (2001) ague that in principle overshadowing is possible in visual learning paradigms with *Drosophila*. If overshadowing occurred in our paradigm, then the faster learning of the single bar may have partially blocked the slower learning of the grating categories.

We observed differences in learning between the bees trained with the CS+ containing the vertical single element and the horizontal single element. Ours is not the first to report bees learn visual and horizontal stimuli at different rates (MaBouDi et al. 2021, Guiraud et al. 2023). Why this may be not clear, but our data are consistent with bees learning the vertical bar as rewarded faster than the horizontal bar as rewarded. This has been seen in other studies (Srinivasan et al. 1999, Wang, Tie et al. 2014, Wolf et al. 2015, Guiraud et al. 2023).

Our experiment shows that bees can rapidly learn multicomponent visual stimuli. Our data are consistent with bees “attending to” the simplest and most consistent element of a multicomponent stimulus, but we do not need to invoke attentional processes to explain our data. Generalisation and overshadowing phenomena - both consequences of Rescorla-Wagner models of learning (Rescola et al. 1972) – are sufficient to explain our data. Our visual stimuli were complex in the computer vision sense of being composed of multiple elements and differing in multiple ways, but we do not need to invoke complex cognitive processes to explain effective and efficient learning of these stimuli.

## Acknowledgements

We thank the London Bee lab for helpful discussions throughout the study. We thank Prof. Lars Chittka for PhD supervision, help with the conception and revision of the manuscript. We thank Aqsa Muhammad (Nuffield) for help with the data collection. We thank Prof. Andrew Barron for his help with the manuscript and his supervision.

## Author contributions

MGG and HM conceived the study. MGG designed the protocol. MGG and EQK acquired the data. MGG curated the data. MGG performed video analysis. VG created the software for video analysis. MGG and HM statistically analysed the data. MGG drafted the manuscript. MGG and HM revised the manuscript.

## Data availability

Data will be provided upon request.

## Ethical approval

Our research involved bumblebees from commercially available colonies dedicated to research for which an approval of an ethical committee is not mandatory. The protocols comply with standard welfare practice in our field and a minimum number of individuals were used to study our scientific question. The animals were not harmed during the experimental procedures and went on to live a happy retired life after experiments.

## Supplementary material

**Supplementary Figure 1.**
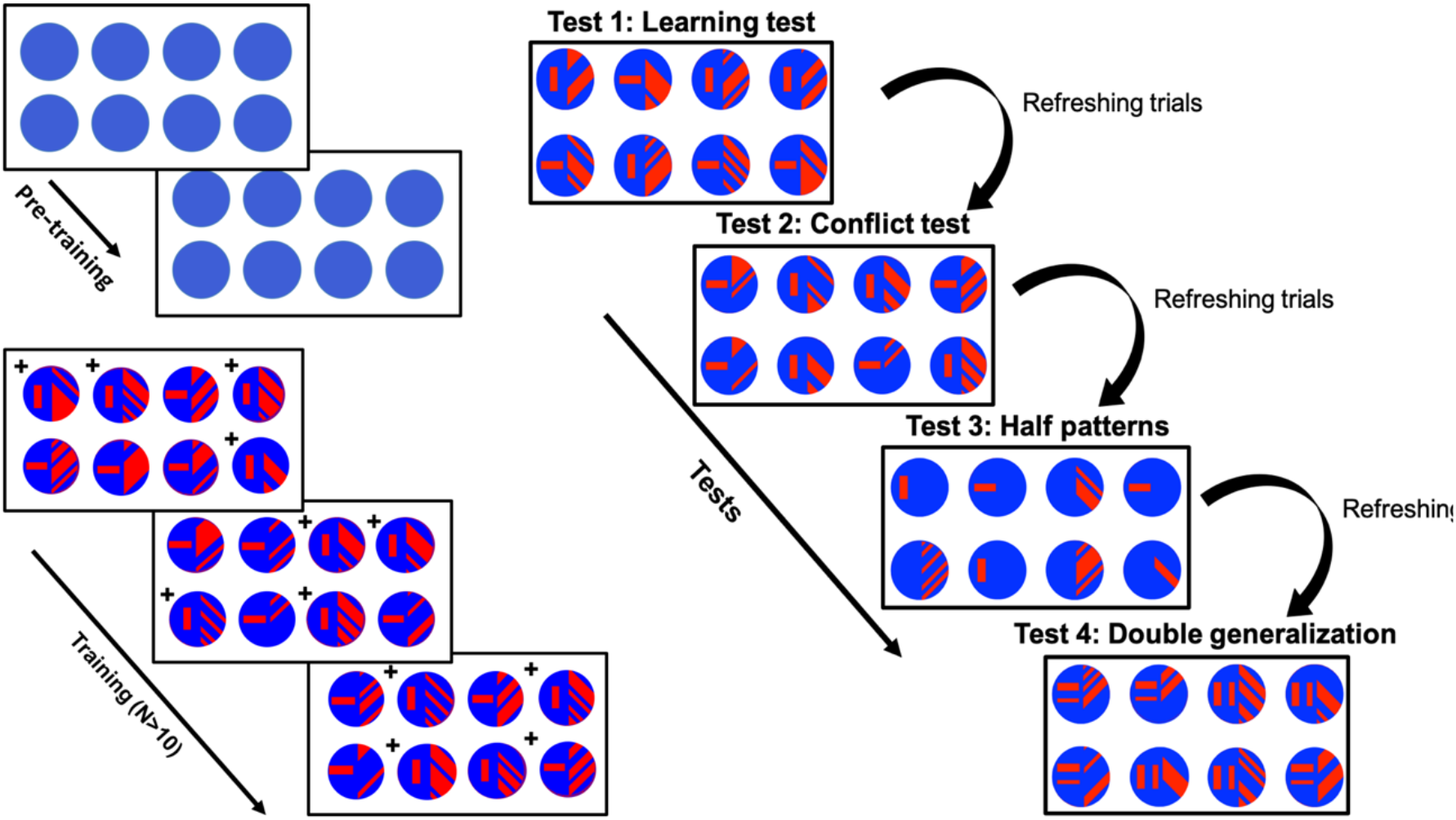
Training and testing protocol. Example of the conditioning and testing procedure. Left panel: bees were exposed to 2-3 pre-training bouts where eight blue circular stimuli were rewarded (50% sugar solution - w/w). Training consisted of trials with four rewarding stimuli (CS+) and four penalized stimuli (providing quinine solution, CS-). Training continued until the bees reached 80% correct choices over 20 consecutive choices. Right panel: Unrewarded tests were subsequently performed with a learning test, conflict pattern test, half-pattern test and generalisation test (see methods for details). The learning test consisted of training patterns the bee wasn’t exposed to. Conflicting test stimuli had the first part (unique bar) of the CS+ stimuli presented with the second part of the CS-(gratings) and *vice versa*. Half-pattern tests: only one part of the stimuli was present (either the unique bar or the gratings from the trained CS+ and CS-configurations). The generalisation test consisted of similar patterns to training ones; the half with the unique bar presented two parallel bars now. Bees’ performance was evaluated based on the number of landings on each presented pattern during 120 sec of the unrewarded tests.

**Supplementary table 1.** Statistical evaluation of the bees’ performance in training and learning test. A. Training. B. Learning test.

| A | Aim of the test | Group of bees | Statistical test | Statistical values |
| --- | --- | --- | --- | --- |
|  | Comparing the proportion of bees' correct choices when reaching criterion with the chance level (50%) | Bees rewarded on LH+45 | One sided Wilcoxon signed rank test | z= 2.51, p=0.0060 |
| | Comparing the proportion of bees' correct choices when reaching criterion with the chance level (50%) | Bees rewarded on RH-45 | One sided Wilcoxon signed rank test | $z = 1.8418$,<br>**$p = 0.0328$** |
| | Comparing the proportion of bees' correct choices when reaching criterion with the chance level (50%) | Bees rewarded on LV+45 | One sided Wilcoxon signed rank test | $z = 2.4870$,<br>**$p = 0.0064$** |
| | Comparing the proportion of bees' correct choices when reaching criterion with the chance level (50%) | Bees rewarded on RV-45 | One sided Wilcoxon signed rank test | $z = 2.4534$,<br>**$p = 0.0071$** |
| | Comparing the length of training<br>Comparing the proportion of bees' correct choices with the chance level (50%) | All four groups | Kruskal-Willis test | $df = 39$, Chi-sq = 2.47,<br>$p = 0.4806$ |
| **B.** | Difference between groups | Bees rewarded on LH+45 | Wilcoxon signed rank test | $z = 2.80$,<br>**$p = 0.0020$** |
| | | Bees rewarded on RH-45 | Wilcoxon signed rank test | $z = 2.81$,<br>**$p = 0.0049$** |
|  |  | Bees rewarded on LV+45 | Wilcoxon signed rank test | z= 2.80, **p=0.0050** |
|  |  | Bees rewarded on RV-45 | Wilcoxon signed rank test | z= 2.93, **p=0.0033** |

**Supplementary table 2.** Statistical evaluation of the bees’ performance in half-pattern tests.

| Aim of the test | Group of bees | Statistical test | Statistical values |
| --- | --- | --- | --- |
| Comparing the responses of bees to the CS+ Single bar **SB+** vs CS- Single bar **SB-** | Bees rewarded on LH+45 | Wilcoxon signed rank test | z= 2.8031, p= 0.0051 |
|  | Bees rewarded on RH-45 | Wilcoxon signed rank test | z= 2.7082, p=0.0068 |
|  | Bees rewarded on LV+45 | Wilcoxon signed rank test | z= 2.6656, p= 0.0077 |
|  | Bees rewarded on RV-45 | Wilcoxon signed rank test | z= 2.8031, p= 0.0051 |
| Comparing the responses of bees to CS+ gratings **G+** vs CS- gratings **G-** | Bees rewarded on LH+45 | Wilcoxon signed rank test | z= 2.4973, p= 0.0125 |
|  | Bees rewarded on RH-45 | Wilcoxon signed rank test | z= 2.3664p= 0.0180 |
| | Bees rewarded on LV+45 | Wilcoxon signed rank test | $z = 0.3557, p = 0.7220$ |
| | Bees rewarded on RV-45 | Wilcoxon signed rank test | $z = 1.8204, p = 0.0687$ |
| Comparing the responses of bees to **SB+** vs **G+** | Bees rewarded on LH+45 | Wilcoxon signed rank test | $z = 1.1220, p = 0.2619$ |
| | Bees rewarded on RH-45 | Wilcoxon signed rank test | $z = 1.6362, p = 0.1018$ |
| | Bees rewarded on LV+45 | Wilcoxon signed rank test | $z = 2.5236, p = 0.0116$ |
| | Bees rewarded on RV-45 | Wilcoxon signed rank test | $z = 2.2439, p = 0.0248$ |

**Supplementary table 3.** Statistical evaluation of the bees’ performance in conflict test.

| Aim of the test | Group of bees | Statistical test | Statistical values |
| --- | --- | --- | --- |
| Comparing the proportion of bees' responses to SB+G- from chance level (50%) | Bees rewarded on LH+45 | Wilcoxon signed rank test | -0.7645, 0.4446 |
| | Bees rewarded on RH-45 | Wilcoxon signed rank test | $Z = 1.1752, p = 0.2399$ |
| | Bees rewarded on LV+45 | Wilcoxon signed rank test | $Z = 2.4973, p = 0.0125$ |
|  | Bees rewarded on RV-45 | Wilcoxon signed rank test | Z=2.8076, p=0.0050 |
| Difference between the H group bees | H group bees | Wilcoxon rank sum test | z= -1.2875, p=0.1979 |
| Difference between the V group bees | V group bees | Wilcoxon rank sum test | z= -0.8464, p=0.3973 |
| Difference between the H and V groups of bees | All four groups | Wilcoxon rank sum test | Z= -1.5674 , p=0.1170 |

**Supplementary table 4.**
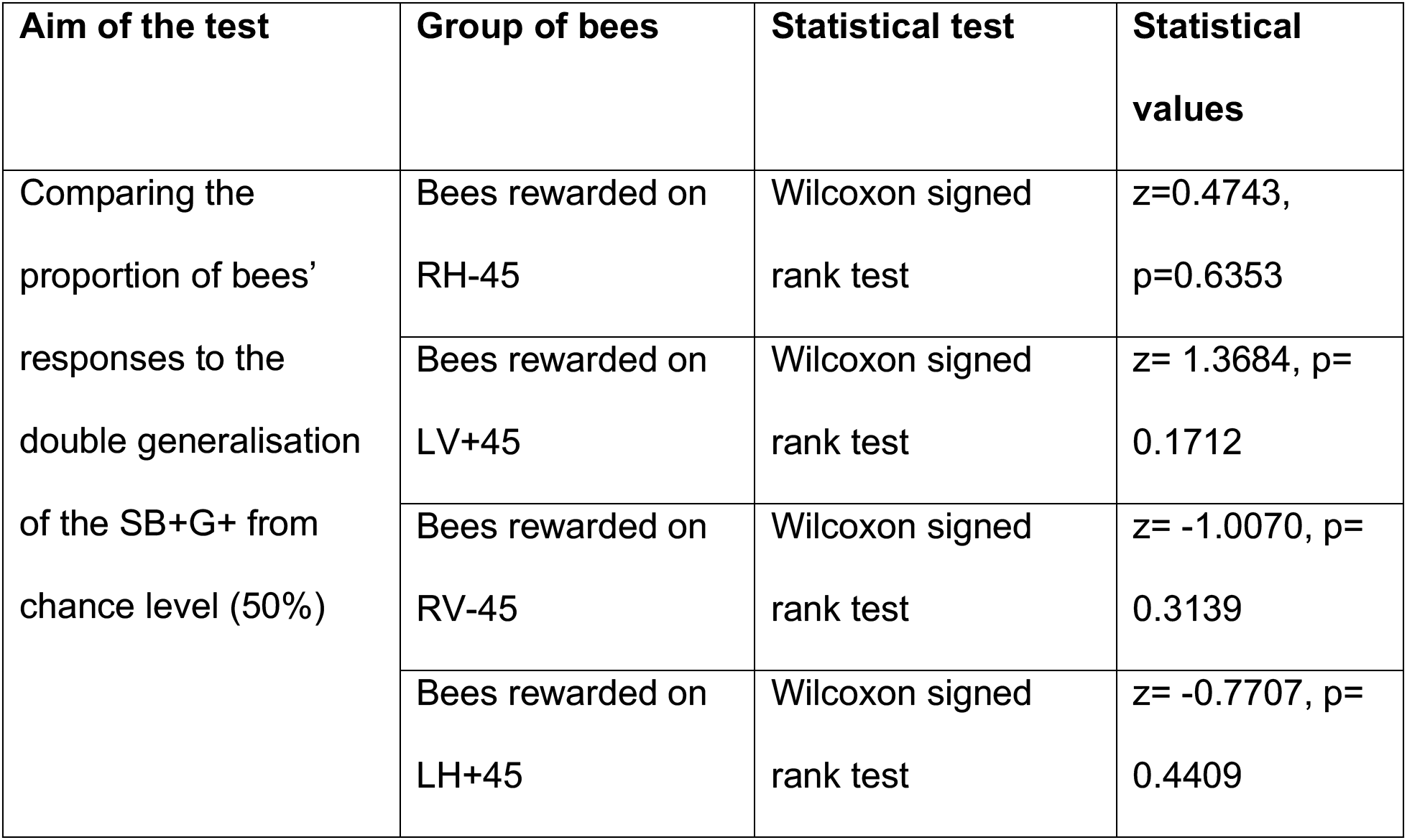
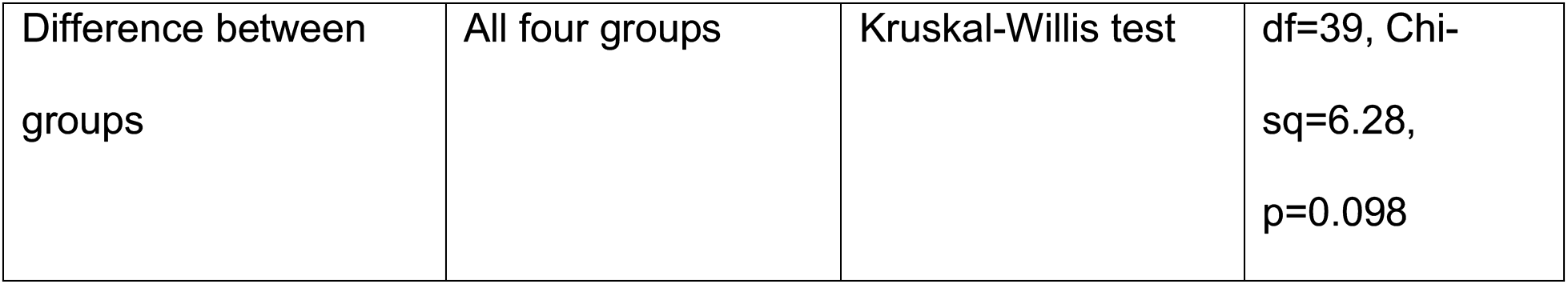
Statistical evaluation of the bees’ performance in generalisation test.

| Aim of the test | Group of bees | Statistical test | Statistical values |
| --- | --- | --- | --- |
| Comparing the proportion of bees' responses to the double generalisation of the SB+G+ from chance level (50%) | Bees rewarded on RH-45 | Wilcoxon signed rank test | z=0.4743, p=0.6353 |
|  | Bees rewarded on LV+45 | Wilcoxon signed rank test | z= 1.3684, p=0.1712 |
|  | Bees rewarded on RV-45 | Wilcoxon signed rank test | z= -1.0070, p=0.3139 |
|  | Bees rewarded on LH+45 | Wilcoxon signed rank test | z= -0.7707, p=0.4409 |
| Difference between groups | All four groups | Kruskal-Willis test | df=39, Chi-sq=6.28, p=0.098 |

